# UBA52 attunes VDAC1-mediated mitochondrial dysfunction and dopaminergic neuronal death

**DOI:** 10.1101/2022.09.22.508987

**Authors:** Shubhangini Tiwari, Abhishek Singh, Parul Gupta, Amrutha K, Sarika Singh

## Abstract

Mitochondrial homeostasis regulates energy metabolism, calcium buffering, cell function and apoptosis. The present study has been conducted to investigate the implications of ubiquitin-encoding gene UBA52 in mitochondrial physiology. Transient expression of Myc-UBA52 in neurons significantly inhibited the rotenone-induced increase in reactive oxygen species generation, nitrite level and depleted glutathione level. Mass spectrometric and co-immunoprecipitation data suggested the profound interaction of UBA52 with mitochondrial outer membrane channel protein, VDAC1 in both the wild-type and Myc-α-synuclein overexpressed neuronal cells and in the Parkinson’s disease (PD)-specific substantia nigra and striatal region of the rat brain. *In vitro* ubiquitylation assay revealed that UBA52 participates in the ubiquitylation of VDAC1 through E3 ligase CHIP. Myc-UBA52 overexpression in neurons further improved the mitochondrial functionality and cell viability by preventing the alteration in mitochondrial membrane potential, mitochondrial complex-I activity, translocation of cytochrome-c and p-Nrf2 along with effect on intracellular calcium uptake, thus collectively inhibiting the opening of mitochondrial permeability transition pore. Additionally, Myc-UBA52 expression in neuronal cells offered protection against apoptotic and autophagic cell death. Altogether, our findings delineate functional association between UBA52 and mitochondrial homeostasis, providing new insights into the deterrence of dopaminergic cell death during acute PD pathogenesis.

## Introduction

Mitochondria are the vital source of energy required to meet all the metabolic demands of the cell and parallelly contribute in various cellular signalling networks that fine-tune the balance between cell survival and death. Dysfunction of mitochondria and oxidative stress have been implicated as primary biochemical hallmarks in Parkinson’s disease (PD) [1–3]. Nigral dopaminergic neurons are highly susceptible to mitochondrial damage as they contain high levels of iron and lipids and increased dopamine metabolism, leading to higher production of reactive oxygen species (ROS). Emerging evidences showed upregulated level of 4-hydroxyl-2-nonenal (HNE)-a lipid peroxidation product in the post-mortem brain of PD patients that cause oxidative stress and neuronal apoptosis in the dopaminergic neurons of substantia nigra [4]. The exposure to neurotoxins such as methyl-4-phenyl-1,2,3,6 tetrahydropyridine (MPTP), rotenone, paraquat-induced experimental model of PD mimic the diseased conditions through inhibition of mitochondrial complex-I (NADH-ubiquinone reductase) activity which further obstructs oxidative phosphorylation (OXPHOS) pathway and ATP production [3,5]. PD-associated genes such as α-synuclein, LRRK2, DJ1, PINK1 and Parkin potentially interact with the mitochondrial proteins and modulate its functionality along with reduced synaptic plasticity and dopamine release [6]. The interaction of α-synuclein, chaperone HSP90 and ubiquitin in the mitochondria has been reported which contribute in Lewy bodies formation and neuronal death during PD conditions [7].

Mitochondrial functions are critically regulated by the channels located on the inner and outer mitochondrial membrane and their opening and closing state allow the fine-tuning of mitochondrial membrane potential, ROS generation and activity of respiratory complexes [8]. One of mitochondrial outer membrane located channel is voltage dependant ion channel (VDAC) which account for approximately fifty percent of the overall protein content and responsible for approximately ninety percent of outer mitochondrial membrane permeability [9,10]. VDAC1 is the highly expressed isoform and overcome the expression of VDAC2 and VDAC3 by one and two orders of magnitude respectively [11] and facilitate the cross-talk between mitochondria and the rest of the cell. In addition to regulation of mitochondrial metabolism and energetic functions, VDAC also appears to be convergence point for a variety of cell survival and death signals [12]. VDAC1 also represents the main mitochondrial docking site of various disease related misfolded proteins, like amyloid β and Tau involved in Alzheimer’s disease, α-synuclein involved in Parkinson’s disease and several SOD1 mutants involved in Amyotrophic Lateral Sclerosis [13].

Activation and opening of VDAC1 channel take place at depolarising potential (−40mV-+40mV) in which it allows the passage of key metabolites such as ATP, organic anions and various respiratory substrates, whereas in the closed state, the transport of Na^+^, K^+^ and Ca^2+^ takes place [14]. In addition to interaction with disease related proteins VDAC1 also interacts with various pro- and anti-apoptotic signalling proteins like Bcl2, Bax, hexokinase and cytosolic kinases [15]. Parallelly, VDAC1 regulates the mitochondrial transition permeability pore (mPTP) opening [16,17], that lead to release of cytochrome-c, ROS generation and initiation of caspase-9 dependent apoptosis [18]. Various research groups have debated upon the dominance of its closed or open state in induction of apoptosis, however, findings suggest that closed conformation of VDAC1 allows influx of Ca^2+^ in the mitochondria which cause Ca^2+^ overload, accumulation of superoxide anions and finally facilitating the Ca^2+^ dependent opening of the mPTP [19]. Ca^2+^ is released from the endoplasmic reticulum (ER) and is taken up by the mitochondria through VDAC1 within the mitochondrial-associated membranes (MAMs) that mediate apoptotic signalling from the ER to the mitochondria [20,21]. In agreement to the above-mentioned study, it has been reported that overexpression of a small non-coding microRNA-7 reduced the expression of VDAC1 and inhibited the mPTP opening [22]. Interaction of hexokinase with VDAC1 prevents its interaction with Bax or Bak, reduces the ROS generation and inhibits apoptosis [23]. During PD, high interaction of α-synuclein with VDAC1 has been shown by [24], wherein VDAC1 channel act as a pathway for α-synuclein entrance into the intermembrane space of the mitochondria and target the mitochondrial membrane located complex I-IV [25–27]. Reports also suggested that α-synuclein severely impacts the mitochondrial functionality and enhance the oxidative stress through alteration of VDAC1 permeability.

Besides, mitochondrial dynamics is also regulated by ubiquitin-linked degradation of proteins either targeted to the proteasome or endosomal-lysosome pathway to prevent their accumulation and restore the mitochondrial quality control [28]. Ubiquitylation of various OMM proteins such as VDAC1, Bax, DRP1, MFN1/2 has been reported by several E3 ligases including Parkin, SIAH2, MARCH5 which have shown their implications in progression of PD pathogenesis (Lavie et al, 2018; Kocaturk and Gozuacik, 2018). Key role of ubiquitin-proteasome system (UPS) in modulating the mitochondrial physiology through interaction and ubiquitylation of various intramembrane space and IMM proteins such as succinate dehydrogenase subunit A (SCHA) and other TCA-cycle metabolites has also been reported [28,29]. Though reports are available showing the link between UPS and mitochondrial dysfunction however, no study reports the participation of ubiquitin encoding genes (UBB; UBC; UBA52; RPS27a) in the mitochondrial functions that are essentially needed to attach ubiquitin residue(s) to the target substrate in UPS.

Recently, we reported the CHIP-UBA52 (Ubiquitin-60S ribosomal protein L40) mediated ubiquitylation of chaperone HSP90 that inhibits ER stress and prevents the altered level of PD-specific pathological markers such as tyrosine hydroxylase (TH) and α-synuclein [30]. We have also found changes in protein level of mitochondria-associated HSP90-client protein PINK1 and HSP75 (homologue of HSP90 in mitochondria) upon UBA52 overexpression, which led us to speculate that UBA52 might show a significant role in altering oxidative stress, mitochondrial dynamics and reducing cell death through interaction with various apoptosis-related proteins. In coherence, here in this study we unravelled the novel role of UBA52 in mitochondria-associated biochemical alterations employing Myc-UBA52 transient overexpression strategy.

## Material and Methods

### Cell Culture and treatments

SH-SY5Y neuroblastoma cell line was obtained from NCCS (National Centre of Cell Sciences), Pune, India. The cells were maintained using a mixture of DMEM/F12 (1:1) tissue culture media, penicillin/streptomycin (GIBCO) and 10% FBS in an atmosphere of 5% CO2 at 37^°^C. Rotenone was dissolved in DMSO and added to the media at 500nM concentration for 24h to induce the PD pathology in cells, validated through PD-specific markers [5]. MG132 (proteasome inhibitor) was added to media at 5μM for 6h before the initiation of respective experiments to inhibit the endogenous proteasome activity. All experiments were repeated at least three times (represented as ‘nexp’) for statistical significance.

### Plasmids, cloning and transfection

Plasmid constructs overexpressing Myc-ddk tagged wild-type human α-syn (Myc-α-synuclein) and Myc-ddk tagged wild-type human UBA52 (Myc-UBA52) were purchased from Origene technologies. Each plasmid was first propagated in DH5α cells and later isolated utilizing the plasmid isolation kit from Qiagen. SH-SY5Y cells were transiently overexpressed with respective plasmids via standard protocol provided for Lipofectamine3000 mediated transfection. Confirmation of transfection was done through immunoblotting using anti-c-myc antibody alongside an empty vector, PcDNA3.1 used as a negative control.

### Animals and stereotaxy

Adult male SD (Sprague-Dawley) rats (200-220g) were employed for the study. Neurotoxin (rotenone) was administered in the SN and STR region of the rat brain to induce dopaminergic neurotoxicity. The animals were routinely maintained in the National Laboratory Animal Centre of CSIR-Central Drug Research Institute. Animal maintenance and all procedures were performed in accordance with ARRIVE guidelines provided by the institutional animal ethics committee [CSIR-CDRI (IAEC/2018/F-52)]. For each set of experiment, rats were divided in a group of two: Sham-control (DMSO-injected) and rotenone-administered. A mixture of xylazine (10mg/kg) and ketamine (80mg/kg) was used to anesthetize experimental rats. Substantia nigra (SN) (5.5mm posterior, 1.5mm lateral, and 8.3mm ventral) and striata (STR) (0.8mm anterior, 2.7mm lateral and 4.5 ventral) brain region was measured from the bregma after mounting the rats on stereotaxic apparatus and rotenone (dissolved in DMSO) (6μg in each region) was administered to induce dopaminergic neuronal damage. The rats were provided appropriate care with consistent application of neomycin& betadine on the head stitches. Three days after stereotaxic surgery, the rats were anesthetized as described previously [5,31]. The midbrain and cortical tissue from whole brain was dissected to isolate SN and STR region, respectively for various experimental studies. All experiments were conducted utilising a minimum of 4 rats per group (represented as ‘nrats’) in an individual set of experiment and for each experiment, at least three replicates were evaluated (represented a ‘nexp’) for statistical analysis.

### MTT Assay

After transient expression of Myc-UBA52 in SH-SY5Y cells, mitochondrial dehydrogenase activity assay (MTT) was performed to estimate the cell viability with or without rotenone treatment as reported previously (Gupta et al, 2019). The absorbance was read using a spectrophotometer at 550nm (Gen5, BioTek) and optical density mean+SEM was illustrated in a graph.

### Protein extraction and western blot

Protein extraction, estimation and immunoblotting with respective antibodies was performed using protocol by [32]. β-Actin was used as a loading control. For cytosolic, nuclear and mitochondrial fractions, the protocol was taken from [33]. After completion, the immunoblots were incubated with Femto Lucent plus HRP substrate (G-Biosciences) and visualized under ChemiDoc XRS+ (Biorad). Mean intensity of individual band of each protein was analysed using ImageJ software (NIH, USA).

### Co-immunoprecipitation (Co-IP)

Protein extraction, estimation of cell and rat tissue lysates was performed, as previously mentioned by [34]. For CoIP, 1-2mg lysate from each sample was incubated overnight at 4°C with anti-UBA52-Protein A-Sepharose bead complex or anti-CHIP-Protein G-Sepharose beads [30]. Non-specific proteins bound to the complex were removed through 4-5 times washing with chilled PBS, followed by boiling of samples to denature the attached proteins at 95°C for 10 minutes using 2X laemmli buffer. Separated proteins were first resolved on 4-20% gradient SDS-PAGE gel and later immunoblotted with respective antibodies for assessing the protein level using chemiluminescence or otherwise the protein bands were excised for mass spectrometric sample preparation and analysis.

### Mass spectrometry

The protocol for in-gel digestion for mass spectrometry used in this study was previously mentioned by [35]. Each sample was mixed with α-CHCA (α-cyano-4-hydroxycinnamic acid) matrix (1:1) and spotted on a MALDI plate to be processed using AB Sciex 4800 MALDI-TOF/TOF mass spectrometer. Recordings were made for all positive ion spectra over m/z 800-4000 Da to assess UBA52 interacting proteins. Maximum of twenty precursors with minimum signal: noise ratio was selected from each spectrum for MS/MS analysis and identified using Protein Pilot (AB Sciex). The corresponding proteins against each spectrum showing high interaction with UBA52 were observed using MASCOT and NCBInr database.

### *In vitro* Ubiquitylation assay

For *in vitro* ubiquitylation of VDAC1, firstly co-immunoprecipitation was performed using overnight incubation of cell or tisue lysates and anti-UBA52 antibody-protein A-Sepharose beads at 4°C. The immunocomplexes obtained were rinsed 4-5 times with chilled PBS and added to a mixture, comprising of reaction components provided in in-vitro ubiquitylation assay kit (Enzo life sciences). The final mixture was incubated at 37°C for 1h and terminated by adding 2X non-reducing laemmli buffer. The samples were then resolved on 12% SDS-PAGE, subjected to immunoblotting with anti-ubiquitin antibody and anti-VDAC1 antibody and later visualized using chemiluminescence to confirm ubiquitylation of VDAC1.

### Reactive oxygen species (ROS) generation assay

This assay was performed utilising the protocol from [32]. The reaction mixture was incubated for 2h in the dark at 37°C after addition of Dichlorofluorescindiacetate (DCF-DA) dye. The fluorescence was detected using a fluorimeter (Varian Cary Eclipse, USA) at excitation/emission spectra of 485/520nm respectively.

### Assessment of Mitochondrial membrane potential (ΔΨm)

The protocol for assessment of change in mitochondrial membrane potential (ΔΨm) using Rhodamine123 dye was utilized from [32]. The reaction mixture was incubated for 1h in the dark at 37°C after addition of Rhodamine123 dye. The fluorescence was detected using a fluorimeter (Varian Cary Eclipse, USA) at excitation/emission spectra of 508/530nm respectively.

For JC1 live-cell imaging, the cells were rinsed with PBS, followed by incubation in JC1 dye for 30 minutes kept in the CO2 incubator at 37°C. Excess stain was removed after 3-4 times washing with PBS and the images were captured by the inverted microscope at 40x (Nikon Eclipse E 200) and quantitated.

### Reduced-Glutathione (GSH) estimation

The protocol for GSH estimation was taken from [32]. The reaction mixture turned yellow and the absorbance was detected using ELISA plate reader (BIO-TEK instruments) at 412nm. The activity was calculated in μg/mg of protein.

### Nitrite level measurement

The protocol for nitrite estimation was taken from [36]. The reaction mixture was incubated for 20 minutes in the dark at 37°C and the absorbance was obtained using ELISA plate reader (BIO-TEK instruments) at 550nm. The activity was calculated in μmole/mg of protein.

### Lactate dehydrogenase estimation

The cellular cytotoxicity estimation protocol was adapted from [37]. NADH (0.5M) was added to the reaction mixture and the absorbance was detected using ELISA plate reader (BIO-TEK instruments) at 340nm at a time interval of 15 seconds for continuous two minutes. The activity was calculated in activity/min/mg of protein.

### Mitochondrial Complex I Activity

Complex-I activity was performed utilizing the protocol from [38]. A reaction mixture of 200μl was set up and 29μl of 5mM NADH was added, followed by the incubation in the dark at 37°C for 10 minutes. The enzyme kinetic activity was detected using ELISA plate reader (BIO-TEK instruments) at 340nm at a time interval of 15 seconds for continuous three minutes. The activity was calculated in mol/min/mg of protein.

### Immunofluorescence through Confocal Microscopy

The immunofluorescence was performed utilizing the protocol from [39]. Confocal microscopy imaging was done in SH-SY5Y cells adhered to the poly-L-lysine coated coverslips to visualize the protein expression and localization of VDAC1, Cytochrome-c and phospho-NRF2. After staining, the coverslips were mounted on the glass slides containing DAPI-anti fade medium. The slides were then visualized under confocal microscope (Leica) and images were captured in a single z-confocal axis.

### Intracellular alteration in calcium level estimation

The protocol for assessment of intracellular calcium level was utilized from [36]. The reaction mixture was incubated with 1μM Fluo-3-AM dye in the dark at 37°C for 1h. The fluorescence was detected using a fluorimeter (Varian Cary Eclipse, USA) at excitation/emission spectra of 506/530nm respectively.

### Statistical analysis

Graphical data were analysed using one-way analysis of variance (ANOVA) and the difference between control and treated sets was analysed by Newman Keul’s test. For confocal microscopy imaging, Image-J was used to analyse the images. Values were expressed as the mean ± SEM and the p value less than 0.05 was considered as statistically significant.

## Results

### UBA52 attenuates oxidative and nitrosative stress and recuperates mitochondrial membrane potential in rotenone treated dopaminergic cells

To determine whether UBA52 participates in alteration of oxidative stress-linked biochemical parameters during PD pathogenesis, we first transiently expressed Myc-UBA52 in SH-SY5Y cells followed by rotenone treatment. Data showed that transient expression of Myc-UBA52 in SH-SY5Y cells did not cause any alteration in assessed parameters and levels were comparable to wild type control cells. Previous reports have revealed that rotenone is a potent inhibitor of complex-I activity, induces oxidative stress and dopaminergic cell death, therefore, we assessed the mitochondrial complex-I activity in rotenone-induced neuronal cells [40–42]. In agreement to previous findings, rotenone treatment caused significant inhibition of the mitochondrial complex-I activity, which was not observed in rotenone treated Myc-UBA52 overexpressed cells suggesting the potent role of UBA52 in ameliorating complex-I activity in dopaminergic cells (Fig 1a). Simultaneously, we also evaluated the NADH-dependent succinate dehydrogenase complex-II activity in SH-SY5Y cells which is a parameter to test mitochondrial metabolic rate through MTT assay after UBA52 overexpression. Rotenone treatment significantly caused the neuronal toxicity which was prevented in Myc-UBA52 overexpressed cells (Fig 1b). ROS and superoxide production majorly arises at complex-I (nicotinamide adenine dinucleotide dehydrogenase) junction of electron-transport chain located on the inner mitochondrial membrane (IMM). Data showed that exposure to rotenone led to high production of ROS species, increase in lactate-dehydrogenase activity (LDH) and decrease in anti-oxidant glutathione (GSH) levels compared to the control cells, which was significantly inhibited in the rotenone treated Myc-UBA52 overexpressed cells (Fig 1c-e). Additionally, immunofluorescent imaging and quantification also depicted the high ROS generation upon rotenone exposure, which was contrary to Myc-UBA52 expressed cells. In parallel, Myc-UBA52 expression also prevented the rotenone induced increase in nitrite level (Fig 1f-h).

**Figure 1.**
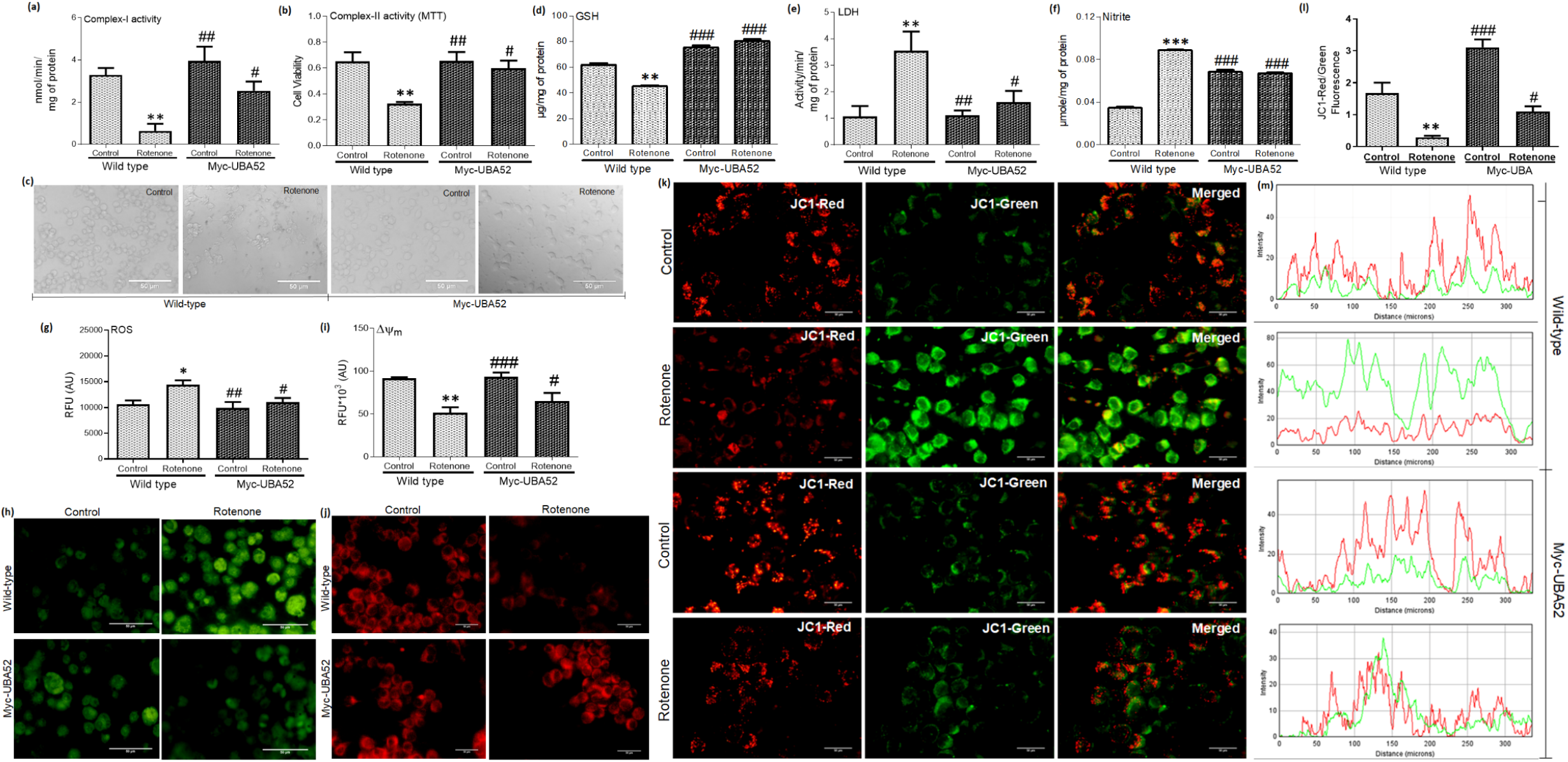
Effect of UBA52 overexpression on oxidative stress markers and altered mitochondrial function In dopaminergic neuronal cells. **(a)** Mitochondrial complex-I activity, **(b)** Cell viability as measured through MTT assay (succinate dehydrogenase complex-II activity), **(c)** Bright-field (20x) images, Scale bar-50μm **(d)** activity Reduced-glutathione (GSH) level **(e)** Lactate dehydrogenase (LDH), **(f)** Nitrite level and **(g&h**) Reactive oxygen species (ROS) generation(Graphical analysis and images) after incubation with DCF-DA dye, Scale bar-50μm; n_exp_=3.In PcDNA3.1 or Myc-UBA52 transfected SH-SY5Y cells exposed to rotenone treatment in comparison to the respective control SH-SY5Y cells. Further, the effect of UBA52 overexpression on depletion in mitochondrial membrane potential in dopaminergic neuronal cell line was assessed. **(i&j)** Representative bar-graph and images (assessed through Rhodamine123 dye) depict the altered mitochondrial membrane potential (Δψ_m_) in PcDNA3.1 (Wild-type) or Myc-UBA52 transfected SH-SY5Y cells with or without rotenone exposure, Scale bar-10μm; n_exp_=3.**(k-m)** lmmunofluorescent images (40x) along with the analysed ratio of JC1-Red/Green fluorescence and the profile plots (depicting the line of fluorescent image pixels as a function of distance) after JC1-staining illustrate the altered Δψ_m_ in PcDNA3.1 or Myc UBA52 transfected SH-SY5Y cells with or without rotenone treatment, Scale bar-50μm; nexp=3.Quantification are mean and SEM of at least three independent experiments and statistical analysis were performed using two-way ANOVA, followed by Tukey’s multiple comparison test. * p<0.05, ** p<0.01, *** p<0.001 control vs. Rotenone; # p<0.05, ## p<0.01, ### p<0.001 Myc-UBA52 vs. Rotenone.

Observed parameters suggested the possible decrease in mitochondrial membrane potential (ΔΨm), therefore we estimated in ΔΨm all the groups. Data suggested the significant decrease in ΔΨm as observed through fluorimeter quantification and imaging, thus facilitating the opening of mPTP to release the pro-apoptotic factors such as cytochrome c into the cytosol to initiate intrinsic apoptotic signalling. To verify the protective role of UBA52, we transiently expressed Myc-UBA52 in SH-SY5Y cells, treated with rotenone and assessed the ΔΨm through Rhodamine123. Rotenone exposure significantly decreased the ΔΨm in comparison to control cells, whereas Myc-UBA52 expression inhibited such alteration. Immunofluorescence imaging through Rhodamine123 and JC-1 staining also simultaneously revealed the similar finding and showed that Myc-UBA52 expressed cells exhibited the inhibition against rotenone-induced depleted ΔΨm in comparison to wild-type rotenone treated cells (Fig i-j). JC1 live imaging in SH-SY5Y cells showed a visible shift from red-stained hyperpolarized mitochondria (negative potential to -140mV) to the green-stain depolarized mitochondria (positive to -100mV) upon exposure to rotenone, which was inhibited in Myc-UBA52 expressing cells (Fig k-m).

Taken together, our results demonstrate that UBA52 plays an important role in ameliorating rotenone induced oxidative stress, mitochondrial dysfunction and nitrite level in dopaminergic neurons.

### UBA52 interacts with and ubiquitylate VDAC1 via E3 ligase CHIP and E2 enzyme UbcH5C

UBA52 attaches ubiquitin residue(s) on the target protein and facilitate ubiquitylation in the presence of E2 and E3 ligases. Reports suggested the inevitable link between UPS and ER homeostasis, which gets compromised during disease pathogenesis, leading to dopaminergic neuronal death [43]. In concordance, our previous study showed neuroprotective role of UBA52 in PD-related ER stress mechanisms [30]. ER-mitochondria cross-talk is also a well-known mechanism that is linked to apoptosis [17]. Besides, we reported that UBA52 overexpression altered the expression of important mitochondrial proteins such as PINK1 and HSP75. Therefore, co-immunoprecipitation studies were performed in both neuronal cells and SN and STR brain region of the SD rat using anti-UBA52 antibody and IgG control to search for novel interacting mitochondrial proteins using mass-spectrometry. MALDI-TOF (matrix assisted laser absorption ionisation-time of flight) and MASCOT search platform revealed OMM protein VDAC1 as one of the key interacting proteins in the mitochondria amongst the series of hits that were obtained during the analysis (Fig 2a). These findings were confirmed further through co-immunoprecipitation and immunoblot analysis using respective antibodies (Fig 2b, e). Next, we transiently expressed Myc-α-synuclein to mimic PD condition and performed co-immunoprecipitation studies using anti-UBA52 antibody with IgG as control to check the interaction of UBA52 and VDAC1. Immunoblots suggested the visible interaction of UBA52 and VDAC1 in both control and Myc-α-synuclein expressed cells. We also observed increased level of VDAC1 upon α-synuclein overexpression, which was coherent with our experimental findings (Fig 2h-j).

**Figure 2:**
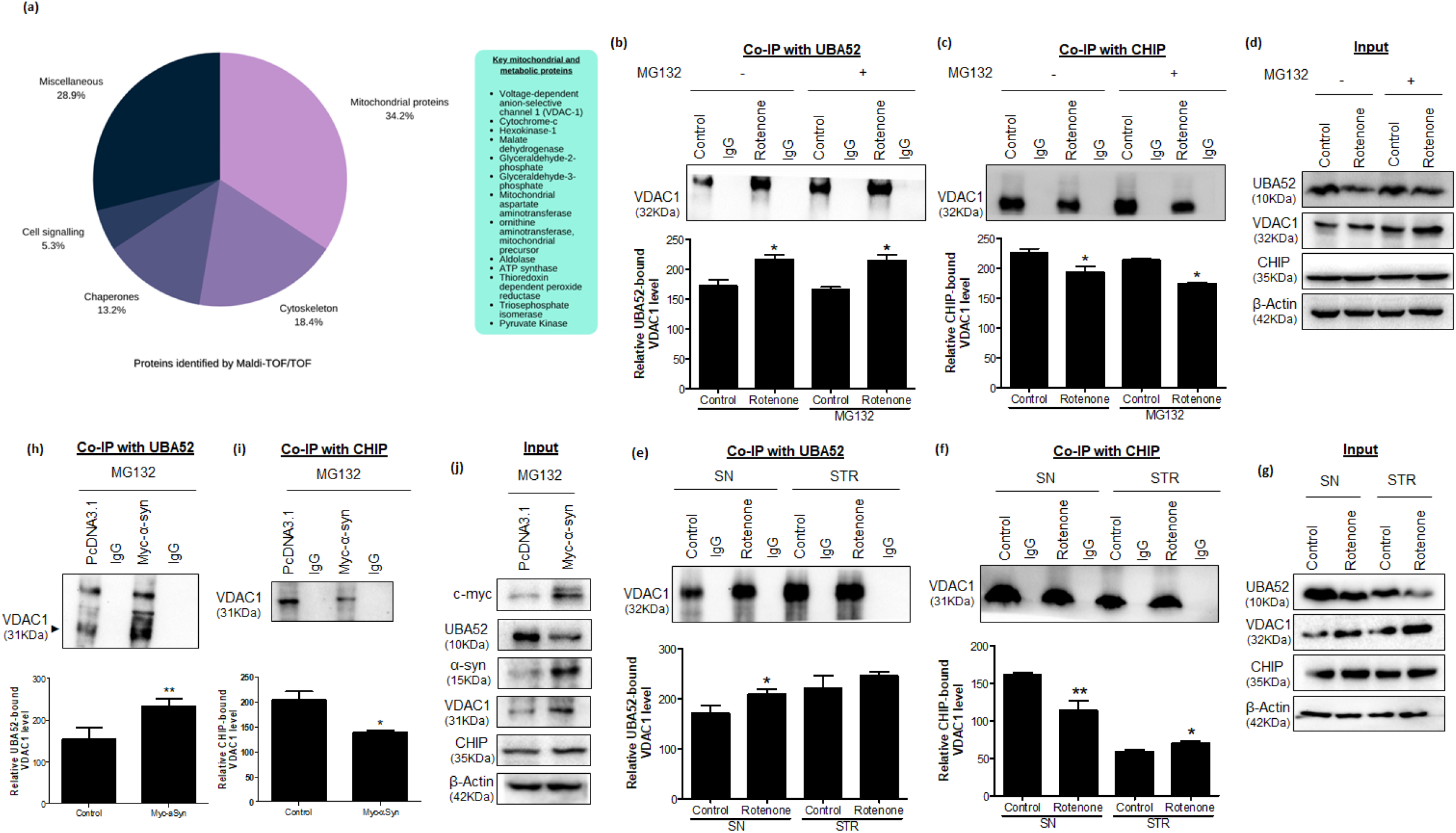
CHIP and UBA52 interacts with the mitochondrial protein VDAC1. Lysates were prepared from the SH-SY5Y cells and the substantia nigra (SN) & striatal (STR) dopaminergic regions of the rat brain and co-immunoprecipitation (Co-IP)-based pulldown of bait proteins was performed using anti-UBA52 or anti-CHIP antibody to assess the interacting proteins of UBA52 and CHIP in both *in vitro* and *in vivo* studies through Mass spectrometry and immunoblotting. **(a)** Pie diagram represents the MS/MS data obtained from the fragmentation spectrum (highest intensity) of the tryptic peptides of various proteins which interact with UBA52 in the SH-SY5Y cells and SN & STR region of the rat brain, including the key mitochondrial proteins such as VDAC1; n_exp_=3. **(b-g)** Representative images and analysed graphs illustrate the Co-IP of UBA52 or CHIP with VDAC1, against lgG as negative control with or without rotenone & MG132 treatment in SH-SY5Y cells; n_exp_=3 and in control and rotenone-lesioned SN and STR rat brain region; total animals used= 9-10 per group; n_exp_=3 along with their respective input bands. **(h-j)** Further, the SH-SY5Y cells were transiently overexpressed with Myc-α-synuclein (a-syn) to mimic the diseased condition and the interaction of UBA52 or CHIP with VDAC1 was assessed against lgG as negative control in PcDNA3.1 or Myc-tagged a-syn transfected SH-SY5Y cells after MG132 treatment; n_exp_ =3. Quantification are mean and SEM of at least three independent experiments and statistical analysis were performed using unpaired t-test (for *in vivo* data) two-way ANOVA, followed by Tukey’s multiple comparison test (for *in vitro*data). * p<0.05, ** p<0.01 control vs Rotenone/Myc-α-Syn.

We further checked the interaction of VDAC1 with few crucial E3 ligases which have already shown their implication in mitochondrial functionality. These E3 ligases were PIAS1, PELLINO1/2, TRAF6, SIAH1/2 and CHIP. Co-immunoprecipitation studies and immunoblot analysis revealed visible interaction of VDAC1 with CHIP in both *in vitro* and *in vivo* control as well as rotenone administered SD rat model of PD (Fig 2c, f), however, the interaction of VDAC1 with other E3 ligases observed through Co-IP and immunoblotting was visibly weak (data not shown) and therefore, VDAC1 was selected for further experiments. After observing strong interaction of CHIP-VDAC1 and UBA52 in our experimental study, we further performed ubiquitylation assay since VDAC1 function in both physiological and pathological conditions may have considerable implications in disease pathogenesis. *In vitro* ubiquitylation assay using various E2 ligases and immunoblot analysis suggested that UBA52 ubiquitylated VDAC1 in the presence of CHIP and E2 enzyme UBCH5c (Fig 3a, b). Altogether, our findings suggested the substantial interaction of UBA52 and CHIP with VDAC1 and ubiquitylation of VDAC1 which is essential in maintenance of mitochondrial homeostasis as it plays significant role in mitochondrial flux as well as in α-synuclein-mitochondria interaction and have significant implications in PD pathology [9].

**Figure 3:**
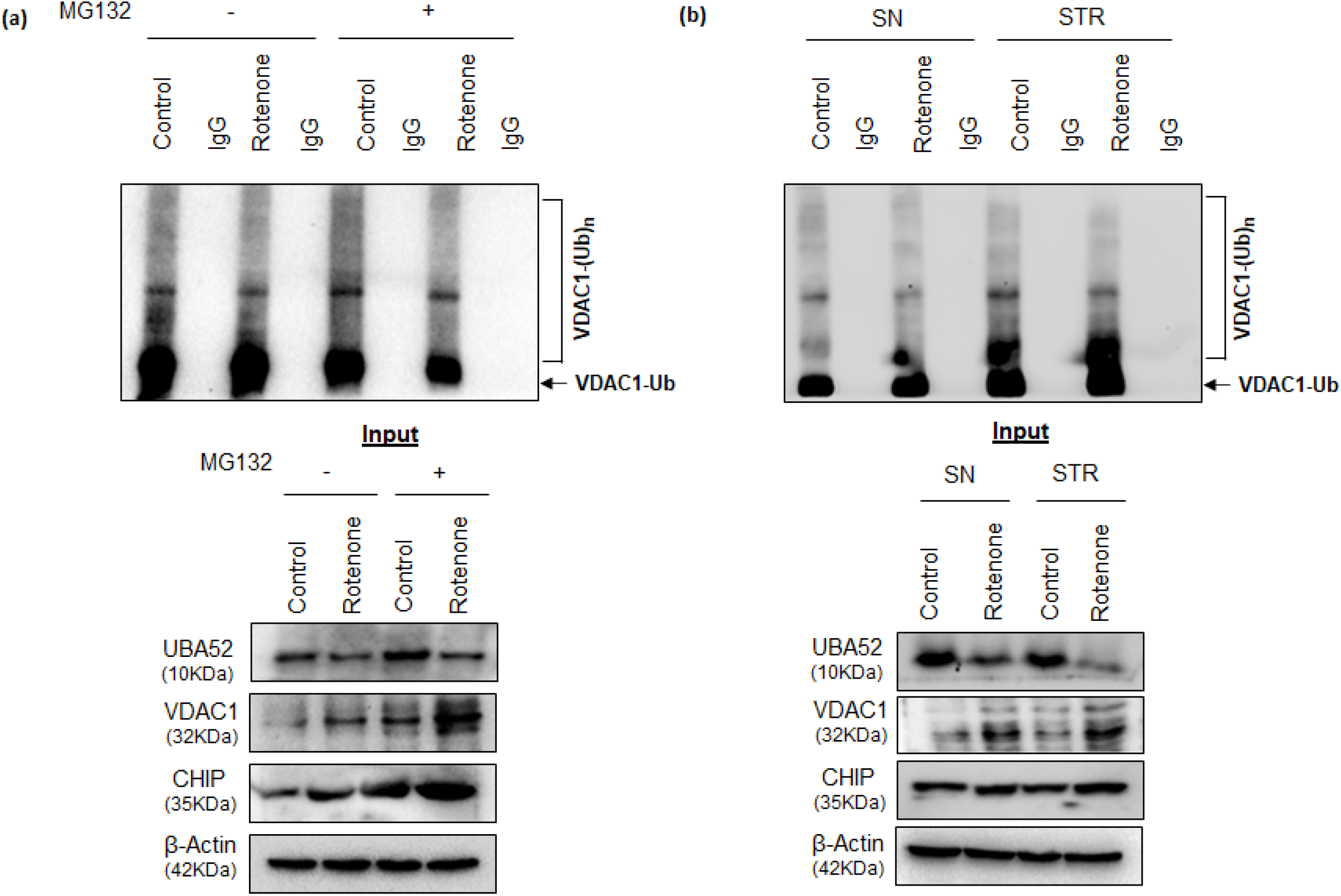
UBA52 mediates the ubiquitylation of VDAC1. Lysates were prepared from the SH-SY5Y cells and the substantia nigra (SN) & striatal (STR) dopaminergic regions of the rat brain and co-immunoprecipitation (Co-lP) based pulldown of bait proteins was performed using anti-UBA52. Further, *in vitro* ubiquitylation assay was set up using anti-UBA52 tagged Protein-A Sepharose beads containing UBA52-interacting proteins and other reaction components, followed by incubation for 90 min at 37°C and the reaction was stopped by adding 2X sample buffer. Final reaction mixture was resolved on 12% gradient SOS-PAGE and immunoblotted using anti-VDAC1 antibody. **(a)** Representative image of *in vitro* ubiquitylation assay in the SH-SY5Y cells after Co-IP of cell lysate with anti UBA52 antibody, against lgG negative control with or without MG132 & rotenone treatment; n_exp_=2 **(b)** lmmunoblots represent the *in vitro* ubiquitylation assay after Co-IP of brain lysate of the control & rotenone lesioned SN and STR region of the rat brain; total animals used= 8-10 per group; n_exp_=2. The positive ubiquitylation reaction was observed in the presence of E2 enzyme, UBCH5c in both the experimental studies.

### UBA52 targets VDAC1 to prevent calcium dyshomeostasis during mitochondrial stress

Since, high interaction of VDAC1 and UBA52 was obtained in co-immunoprecipitation studies, we speculated that UBA52 overexpression would have direct effect on the expression and functional aspect of VDAC1. Interestingly, transient expression of Myc-UBA52 in SH-SY5Y cells inhibited the upregulation of VDAC1 protein level with or without rotenone treatment, in comparison to the wild-type control and rotenone-only treated cells. Next, we also assessed the alteration in VDAC1 expression through immunofluorescence using anti-VDAC1 antibody and mitotracker (to label the mitochondria). In rotenone-treated cells, VDAC1 expression was markedly high in comparison to the control cells and Myc-UBA52 expressed cells. We also observed remarkable reduction in mitotracker expression in rotenone-treated group, which suggested impairment of mitochondria in rotenone-induced toxic conditions (Fig 4b-e).

**Figure 4:**
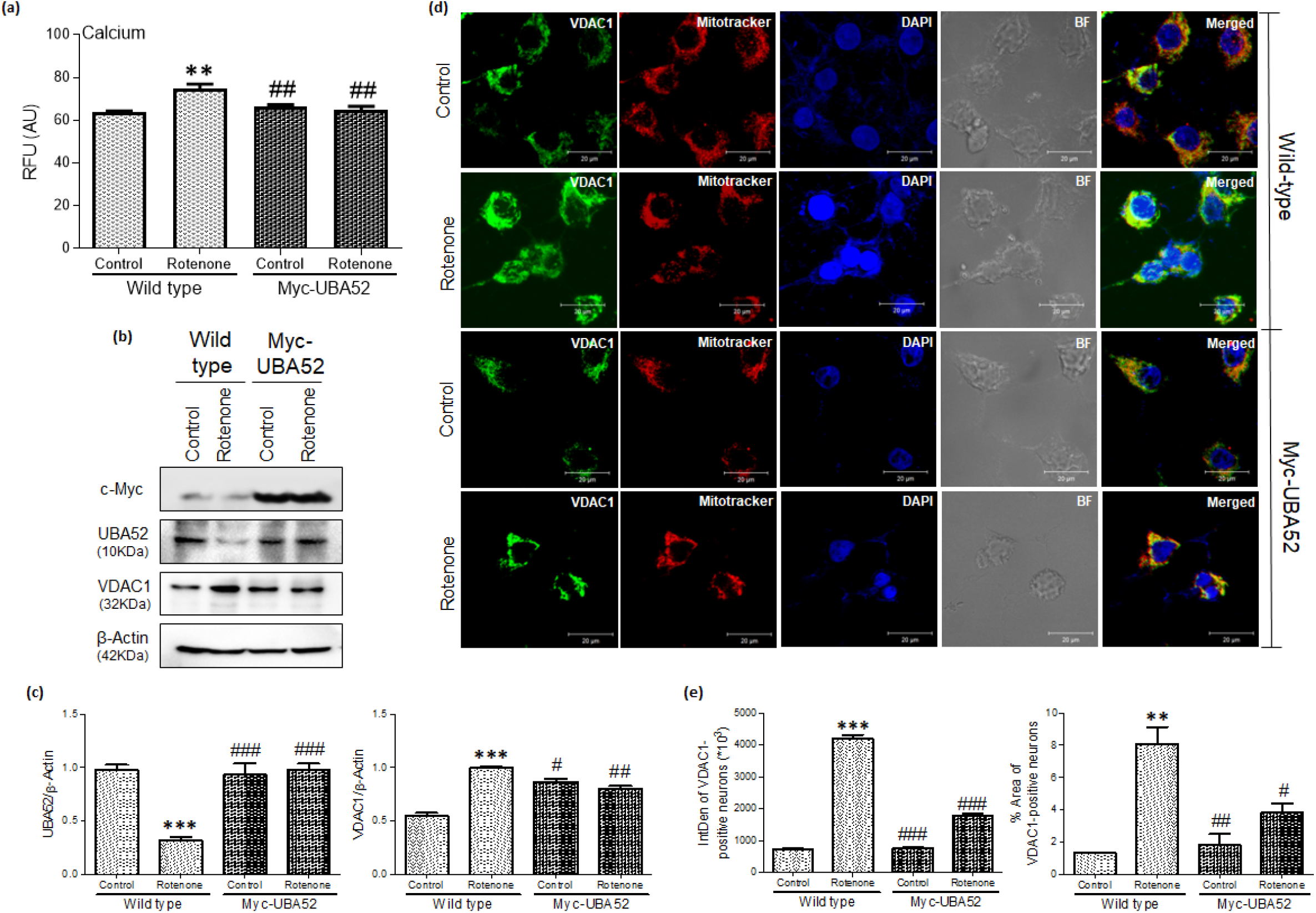
UBA52 regulates the expression of VDAC1 and the linked intracellular calcium influx. **(a)** Graphical representation for intracellular calcium level in PcDNA3.1 (Wild-type) or Myc-UBA52 transfected SH-SYSY cells with or without rotenone treatment; nexp=3. (b&c) Blots and graphs represent the analyzed protein expression of VDAC1 in PcDNA3.1 or Myc-UBA52 transfected SH-SYSY cells with or without rotenone treatment and normalized against β-Actin; n_exp_=3. **(d&e)** Confocal microscopy images (100x) and the integrated density and %Area graph of the VDAC1-positive neurons represent the expression of VDAC1 colocalized with mitotracker (deep red) and counterstaining of DAPI (blue) with or without rotenone in PcDNA3.1 or Myc-UBA52 transfected SH-SYSY cells, Scale bar-20μm; n_exp_ = 3. Quantification are mean and SEM of at least three independent experiments and statistical analysis were performed using two way ANOVA, followed by Tukey’s multiple comparison test. ** p<0.01, *** p<0.001 control vs. Rotenone; ## p<0.01, ### p<0.001 Myc-UBA52 vs. Rotenone.

VDAC1 is the major mitochondrial channel protein that transports various metabolic ions across mitochondria and cytosol, including calcium (Ca^2+^) ions. Increase in mitochondrial calcium either through import from ER or extracellular calcium pool increases mitochondrial damage and facilitate the opening of mPTP. In view of this, we checked the intracellular calcium level of neuronal cells upon UBA52 overexpression. Rotenone treatment significantly upregulated the calcium level in SH-SY5Y cells in comparison to control, which was significantly inhibited in the Myc-UBA52 expressed neurons with or without rotenone treatment (Fig 4a). Our findings showed that UBA52 overexpression participate in the regulation of VDAC1 levels as well as its functional aspect related to intracellular calcium levels, which indeed maintains mitochondrial membrane integrity.

### UBA52 protects dopaminergic cells against mitochondrial-stress induced apoptosis

We observed the rotenone-induced depletion in ΔΨm which may induce the translocation of cytochrome-c from mitochondria to the cytosol, following opening of mPT pore formation. Therefore, further the cytosolic level of cyt-c was estimated in both wild type and Myc-UBA52 overexpressed cells with and without rotenone treatment. The results showed that rotenone treatment caused the translocation of cytochrome-c into the cytosol, which was not observed in Myc-UBA52 expressed SH-SY5Y cells. Immunofluorescent imaging also revealed the considerable translocation of cytochrome-c (Fig 5a-c). Data suggested the significant functional contribution of UBA52 in restoration of mitochondrial functionality by preventing changes in membrane potential and release of apoptosis-inducing cytochrome-c.

**Figure 5:**
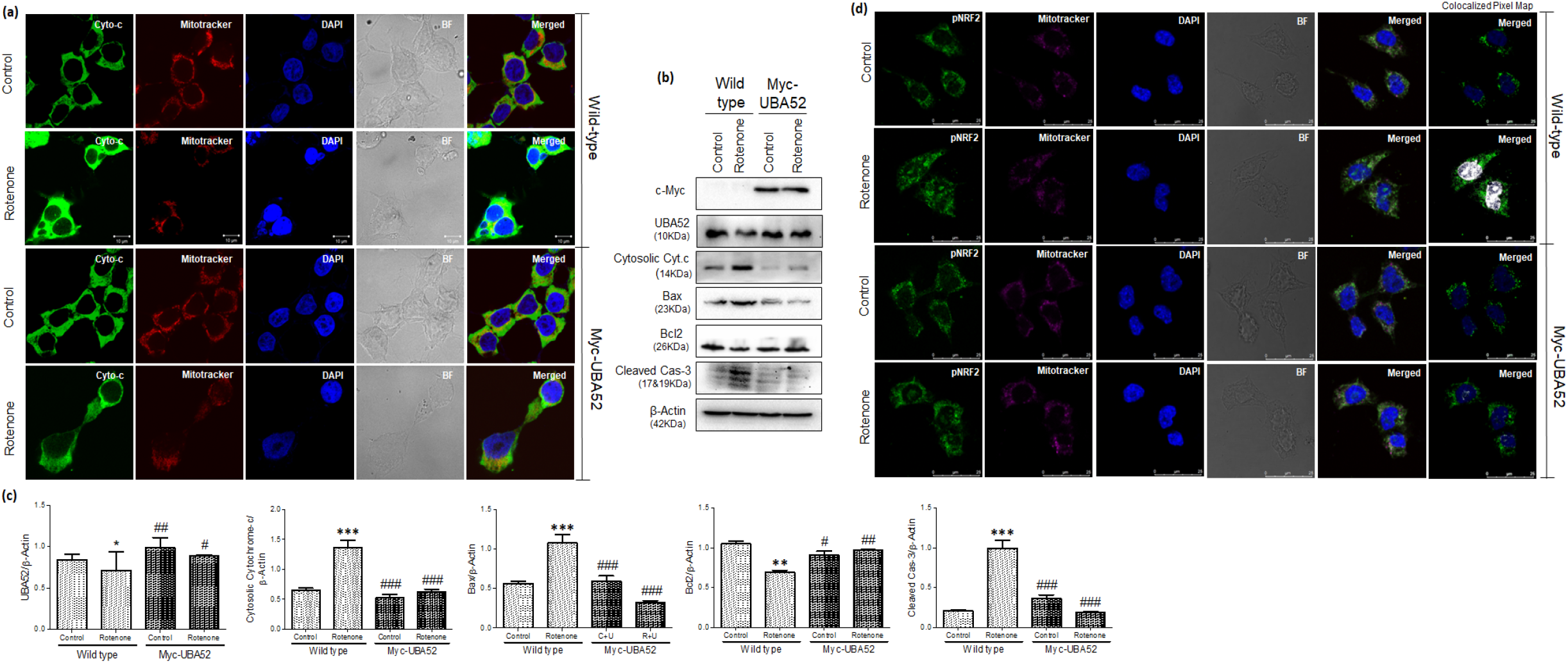
Transient expression of UBA52 inhibits the apoptotic initiation and altered expression of apoplotic markers. **(a)** Confocal microscopy images (100x) represent the expression of cytochrome-c (cyto-c) colocahzed with mitotracker (deep red) and counterstaining of DAPI (blue) with or without rotenone in PcDNA3.1 or Myc-UBA52 transfected SH-SY5Y cells, Scale bar-10μm; n_exp_=3. (**b&c**) Blots and graphs represent the analyzed protein expression of apoptotic markers in PcDNA3.1 or Myc-UBA52 transfected SH-SY5Y cells with or without rotenone treatment and normalized against β-Actin; n_exp_ =3. **(d)** Confocal microscopy images (100x) and the respective colocalized pixel map represent the translocation of the key regulator of oxidative stress phospho-NRF2 from mitochondria (labelled with mitotracker-deep red) to the nucleus (labelled with DAPI) with or without rotenone in PcDNA3.1 or Myc-UBA52 transfected SH-SY5Y cells, Scale bar-25μm; n_exp_ = 3. Quantification are mean and SEM of at least three independent experiments and statistical analysis were performed using two way-ANOVA, followed by Tukey’s multiple comparison test. * p<0.05, ** p<0.001, *** p<0.001 control vs Rotenone; #p<0.05. ## p<0.01, ### p<0.001 Myc-UBA52 vs. Rotenone

It has been reported that cytochrome-c release into the cytosol allows its interaction with various pro- and anti-apoptotic proteins to induce caspase-9 dependent apoptosis [17]. In concordance, we assessed the protein level of Bax, anti-apoptotic Bcl2 and caspase-3 in both wild type and Myc-UBA52 overexpressed cells with and without rotenone treatment. Rotenone treatment significantly caused the alteration in the protein level of Bax, Bcl2 and caspase-3, which was notably prevented in Myc-UBA52 expressed neuronal cells with rotenone treatment (Fig 5b, c).

The role of nuclear factor erythroid 2-related factor-2 (Nrf2) has been strongly implicated in the diseases linked with oxidative stress [44]. During stress conditions, Nrf2 is translocated into the nucleus to regulate its transcriptional machinery, activating anti-oxidant defence signalling and reduces oxidant pathology and apoptosis. Thus, we estimated the phosphorylated-Nrf2 translocation in neuronal cells with or without rotenone treatment in both wild type and Myc-UBA52 overexpressed cells. Confocal microscopy images suggested that rotenone exposure increased the translocation of the phospho-Nrf2 in the nucleus which was attenuated in the Myc-UBA52 transfected neurons and suggest the role of UBA52 in apoptotic signalling (Fig 5d).

### UBA52 surfeit inhibits enhanced autophagy

The interplay between mitochondrial dysfunction and autophagy is also a common mechanism in PD as damaged mitochondrial accumulation demands immediate organelle and its associated protein clearance [45,46]. However, during disease progression, the process is severely compromised, further contributing to disease pathogenesis. In view of this, we checked the protein levels of major autophagy markers such as Beclin-1, LC3-II, Lamp2A and p62/SQSTM-1. Rotenone treatment reduced the protein level of Beclin-1 and Lamp2A as observed through immunoblot analysis, suggesting the reduction in autophagy activation and autophagosome formation. However, transient expression of Myc-UBA52 inhibited such alteration in the respective experimental group. p62/SQSTM-1 and LC3-II accumulation has been observed during autophagy defects and its inhibition restores autophagic activity [47]. To aim this, we checked the protein level of p62/SQSTM-1 and LC3-II in both wild type and Myc-UBA52 expressed cells with or without rotenone treatment. Data showed that Myc-UBA52 overexpressed cells inhibited the upregulation of p62/SQSTM-1 and LC3-II upon rotenone exposure (Fig 6a, b). Findings suggested that UBA52 attenuates the pathologically altered autophagy and contributes in restoration of cellular homeostasis and implicate the significant role of UBA52 in autophagy, disclosing UBA52 participate in regulation of autophagy during dopaminergic neuronal death during PD conditions.

**Figure 6:**
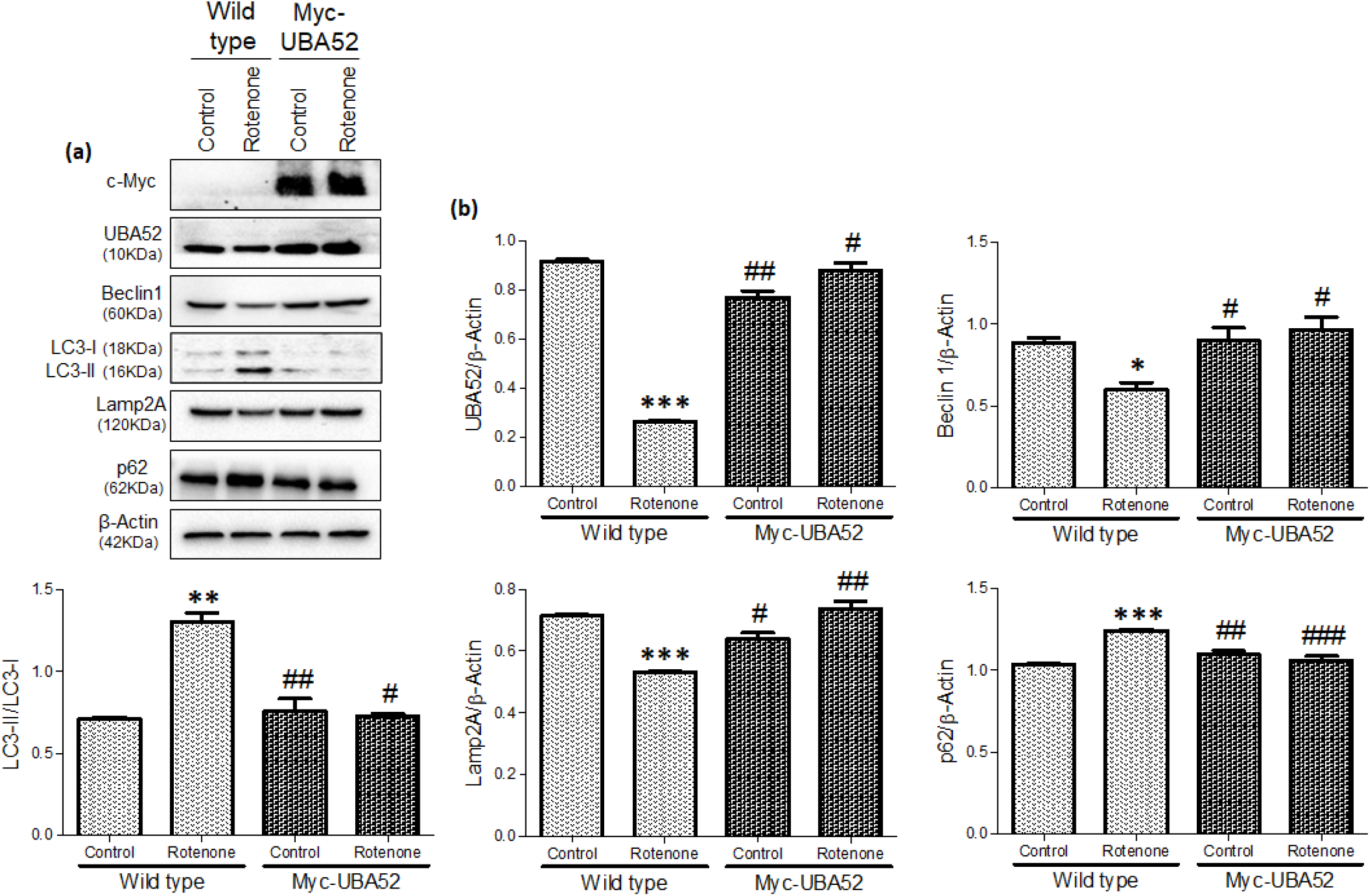
Transient overexpression of UBA52 in SH-SY5Y cells regulates the expression of autophagy marker proteins. **(a)** SH-SY5Y cells were transfected with PcDNA3.1 or Myc-UBA52 with or without rotenone treatment and protein expression of key autophagic markers were assessed through immunoblotting with indicated antibodies. **(b)** Graphs represent the statistical analysis of various proteins normalized against the loading control; n_exp_ =3.Quantification are mean and SEM of at least three independent experiments and statistical analysis were performed using two way-ANOVA, followed by Tukey’s multiple comparison test. * p<0.05, ** p<0.01, *** p<0.001 control vs. Rotenone; # p<0.05, ## p<0.01, ### p<0.001 Myc-UBA52 vs. Rotenone.

## Discussion

Here in this study, we showed the neuroprotective role of UBA52 through its participation in maintenance of redox balance, mitochondrial functions, and neuronal viability. Mitochondrial dysfunction particularly the impairment of complex-I activity has been implicated in PD pathology since nineties [3] and recently been reported to be as therapeutic target for AD [48]. The present study was focused on the pathogenesis of PD-specific dopaminergic neurons employing rotenone induced experimental PD model which is of great interest and mimic most of the PD pathology specific mechanistic events [42,49,50]. Mitochondrial defects are also the key mediators in progression of neurodegenerative disease as ATP is essentially required for protein quality control machinery to restore the cellular homeostasis [1]. The regulation of mitochondrial homeostasis has been reported by some of the ubiquitin E3 ligases as Parkin, MARCH5 and Mul1 [51] however, the studies are still obscure.

Recently, we have reported the significant role of UBA52 in PD pathogenesis [30]. Since mitochondrial dysfunction is one of the critical pathological events in PD pathology the present study was targeted to understand the role of UBA52 in mitochondrial functions in rotenone-induced diseased conditions. We observed that mitochondrial complex I activity and cell viability was significantly reduced in rotenone treated neuronal cells, whereas in Myc-UBA52 transfected cells, it was like control neuronal cells, suggesting that UBA52 may also be involved in OXPHOS pathway and mitochondrial health. ATP generation through OXPHOS leads to ROS generation during physiological conditions which is counterbalanced through various scavenging systems. However, during pathological conditions, enhanced ROS production occurs which imbalanced the internal redox environment and mitochondrial ability to synthesize ATP and participate in other metabolic activities. In accordance, we observed the significantly increased intracellular ROS generation along with reduced GSH level in dopaminergic neurons upon rotenone exposure. Data showed significant downregulation in ROS generation, LDH activity and upregulation in reduced-GSH level after UBA52 overexpression in neuronal cells, showing crucial partaking of UBA52 in sustaining redox balance of dopaminergic neurons. Further, Nrf2 has potentially emerged as the key resistance to oxidative stress by directly targeting ROS and RNS species and participate in the redox-related signalling pathways such as autophagy, ER stress/UPR, mitochondrial homeostasis and apoptotic cell death [44,52,53]. Its activity is regulated by the actin linked-keap1 protein in the cytoplasm, which degrades Nrf2 through UPS pathway in the physiological state and this signalling is disrupted during stress induction, allowing its translocation to the nucleus for transcriptional regulation [54]. Previous findings showed that Nrf2 knockout mice is susceptible to MPTP-induced PD pathology and mitochondrial damage linked with oxidative stress [55,56]. Last year we have also reported the nuclear translocation of Nrf2 in PD conditions [44]. In agreement, here we observed that transient expression of Myc-UBA52 visibly prevented the translocation of Nrf2 in the nucleus in rotenone-treated neurons, thereby, protecting the dopaminergic neurons against stress-activated Nrf2 transcriptional response.

VDAC1 also maintains the crosstalk between the mitochondria and outside through exchange of Ca^2+^ ions and various metabolites [17]. In concordance, further quantitative mass-spectrometric and immunoprecipitation findings showed the high interaction of UBA52 and VDAC1 in both control and pathological conditions. In PD specific rat brain SN and STR regions also high interaction of CHIP and VDAC1 was observed. *In vitro* ubiquitylation data showed that UBA52 participated in the ubiquitylation of VDAC1 in an ATP-dependent manner utilising the E2 enzyme UbcH5c, associated mainly with K48-linked degradative ubiquitylation. Our data suggested that CHIP-UBA52-VDAC1 allied ubiquitylation is a critical signalling component in regulation of mitochondrial-mediated neuronal apoptosis and suggests a protective role of UBA52 in dopaminergic neurons through interaction with VDAC1 and modulating its protein abundance. Besides, transient UBA52 overexpression in neurons inhibited the increase in intracellular calcium flux suggesting its probable participation in calcium level related signalling events. It has been reported that inhibited mitochondrial complex I activity, ROS generation, augmented VDAC1 level and intracellular calcium level collectively leads to loss of mitochondrial membrane potential [57]. Inhibited mitochondrial complex I activity alone could also modulate the mPTP opening to initiate caspase-3-dependent apoptotic signalling [58]. We have observed the non-significant alteration in ΔΨm in the control UBA52 overexpressed cells. In rotenone treated UBA52 transfected cells the depletion in ΔΨm was less in comparison to scramble siRNA transfected rotenone treated neuronal cells suggested that UBA52 exhibit a prominent role in regulating the mitochondrial membrane potential in rotenone-mediated neurotoxicity and may act as a critical interface between mitochondria and cytosol. Studies have also suggested the role of VDAC1 oligomerization and its increased abundance increased mPTP opening propensity which facilitates cytochrome-c translocation [59]. Besides, the interaction of both VDAC1 & cytochrome-c with both pro- and anti-apoptotic proteins such as Bcl2, t-Bid, Bax and Bak has also been reported which further effectuate the opening of mPTP [59]. The mPTP mediated release of apoptotic factors lead to caspase-dependent neuronal apoptosis as reported in neurodegenerative conditions [60,61]. Data showed that transient overexpression of UBA52 in neurons exhibited the reduced translocation of cytochrome-c thus inhibited the mitochondria mediated intrinsic apoptotic event. We have also observed the augmented level of pro-apoptotic and decreased level of anti-apoptotic factors which were reverted to physiological level upon transient expression of Myc-UBA52.

We further suggest that increased mitochondrial permeability enhances mitochondrial damage and fragmentation, initiating autophagy which may be detrimental to the neuronal cell in its inability to replenish healthy mitochondria in replacement to the non-selective mitochondrial clearance [62]. In line with this evidence, our data showed significant alteration in level of autophagy markers in diseased conditions which was inhibited upon transient overexpression of UBA52 though detailed investigation can be done further. Irrespective of the further downstream signalling, our data markedly presents the significant contribution of UBA52 in prevention of apoptosis and autophagic dopaminergic neuronal death. In conclusion, study demonstrates that UBA52 potentially participates in the maintenance of cellular redox environment and mitochondrial longevity through VDAC1 and both apoptosis and autophagy events. However, further preclinical and clinical studies in *in vivo* models will be required for determining the therapeutic efficacy of UBA52 in restoring healthy aging and reducing pathology related to mitochondrial-linked diseases such as Parkinson’s disease.

## Author Contribution

Shubhangini Tiwari and Abhishek Singh conceptualized, designed the work plan and performed the experiments; Parul Gupta and Amrutha K contributed in the experimental work and Shubhangini Tiwari wrote the manuscript; Sarika Singh designed and supervised the study as well as reviewed and edited the final manuscript.

## Acknowledgement

We thank Dr Ramesh Sharma, Ms Anupma Saxena, Ms Kavita Singh, Ms Rima Ray Sarkar and Mr Toofan Raut and SAIF facility (CSIR-CDRI) for technical assistance during the experiments. We further thank Dr Sharad Sharma for auditing our data and providing critical reviews. Web-based tools Biorender and Canva were used for graphical abstract and Pie-chart illustration, respectively. We also acknowledge University Grants Commission (UGC), New Delhi, India for providing one-time research grant and Jawahar Lal Nehru University (JNU) for the research opportunity.

## Ethics approval

Animals were used after approval from the Institutional animal ethics committee of CSIR –

Central Drug Research Institute (IAEC/2018/F-52). For humans: Not applicable.

## Consent for publication

All authors read and approved the final manuscript.

## Availability of data and material

All the generated data is compiled and given in MS or Supplementary file. The raw data will be provided on reasonable request.

---

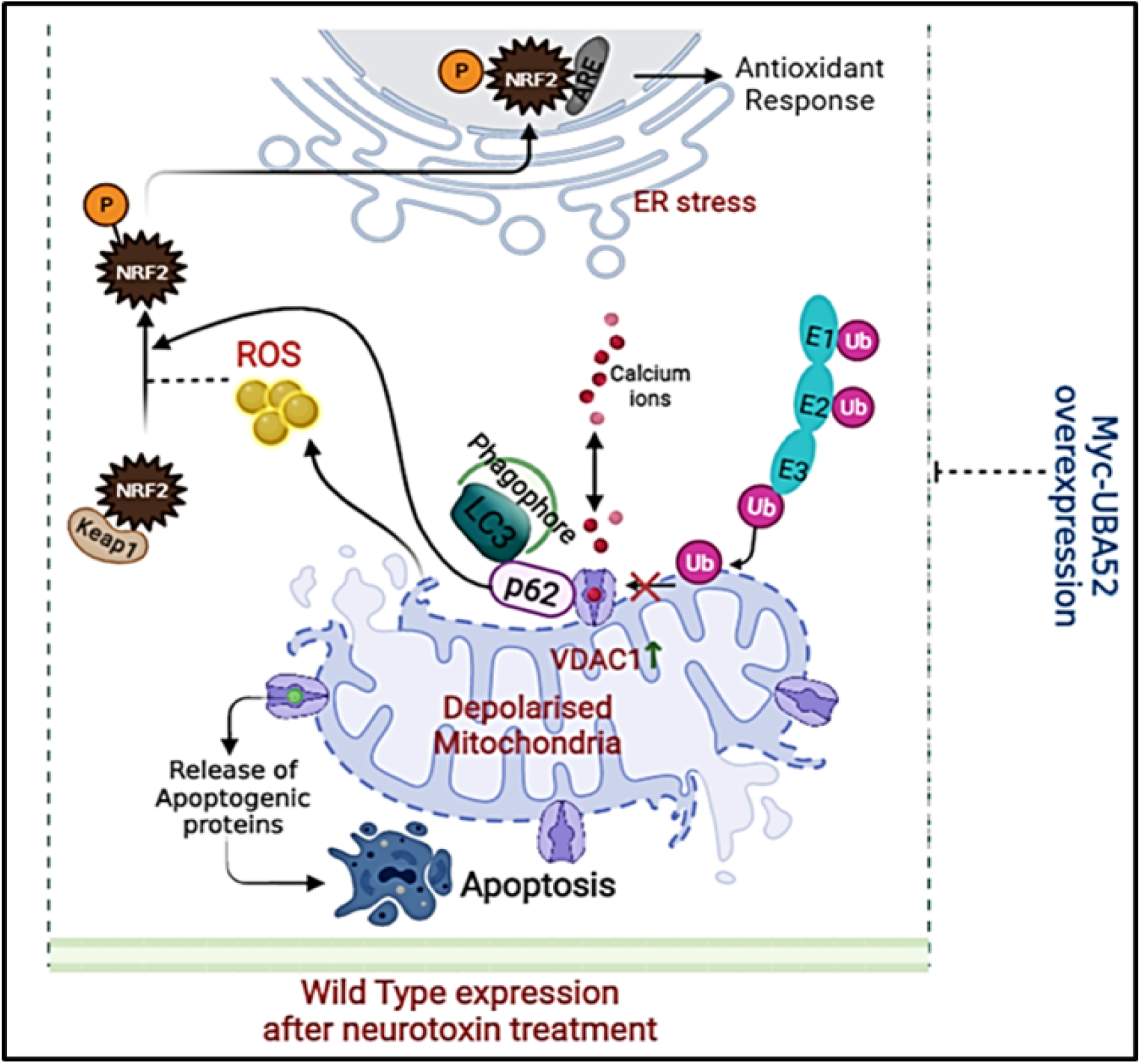

Figure:

UBA52 overexpression prevents the loss of mitochondrial dysfunction and induction of cell death mechanisms. (a) Rotenone, a mitochondrial complex I inhibitor causes oxidative stress and mitochondrial damage, leading to outer mitochondrial membrane (OMM) depolarisation. (b) NRF2 protein abundance is kept low through Keap1-ubiquitin mediated degradation. However, during oxidative stress NRF2 is phosphorylated and dissociated from Keap1 and translocated to the nucleus to activate antioxidant response through interaction with ARE (antioxidant response element). (c) Depolarised mitochondria enhance the interaction of VDAC1 with various apoptogenic proteins and participate in the release of cytochrome-c to the cytosol to initiate apoptosis. (d) Autophagy receptor p62 recognizes the altered VDAC1 conformation and forms complex with LC3 to initiate autophagic signaling.

## Supplementary

### Chemicals

Bovine serum albumin (BSA), disodium hydrogen phosphate (Na2HPO4), dimethyl sulfoxide (DMSO), ethidium bromide, glucose,4-(2-hydroxyethyl)-1-piperazine ethane sulfonic acid (HEPES), NP-40, phenylmethylsulphonyl fluoride (PMSF), sucrose, magnesium chloride (MgCl2), dithiothreitol (DTT), thiobarbituric acid, nicotinamide adenine dinucleotide phosphate reduced (NADPH), sodium bicarbonate, beta mercaptoethanol and tris-buffer were procured from SRL, India. Dulbecco’s modified Eagle’s medium (DMEM), fetal bovine serum (FBS), Trizol, Ham’s F12 medium, penicillin-streptomycin, nuclease-free water, lipofectamine 3000 and mitotracker-deep red were purchased from Invitrogen (San Diego, CA, USA). Luria Bertani broth and agar powder were purchased from Himedia. *In vitro* ubiquitylation kit was purchased from Enzo-Life sciences. Other chemicals such as Protein-A/G Sepharose beads, anti-fade medium DAPI, copper sulfate (CuSO4), calcium chloride (CaCl2), Folin–Ciocalteu reagent, potassium chloride (KCl), sodium carbonate (NaHCO3), sodium chloride (NaCl), sodium dihydrogen phosphate (NaH2PO4), sodium potassium tartrate, sodium pyruvate, sodium hydroxide (NaOH), poly-l-lysine, rhodamine123, protease, nitrite reductase, 5,5’-Dithio-bis(2-nitrobenzoic acid) and phosphatase inhibitor cocktail, rotenone, paraformaldehyde (PFA), ethylenediaminetetraacetic acid (EDTA) were obtained from Sigma, USA.

**Table 1:**
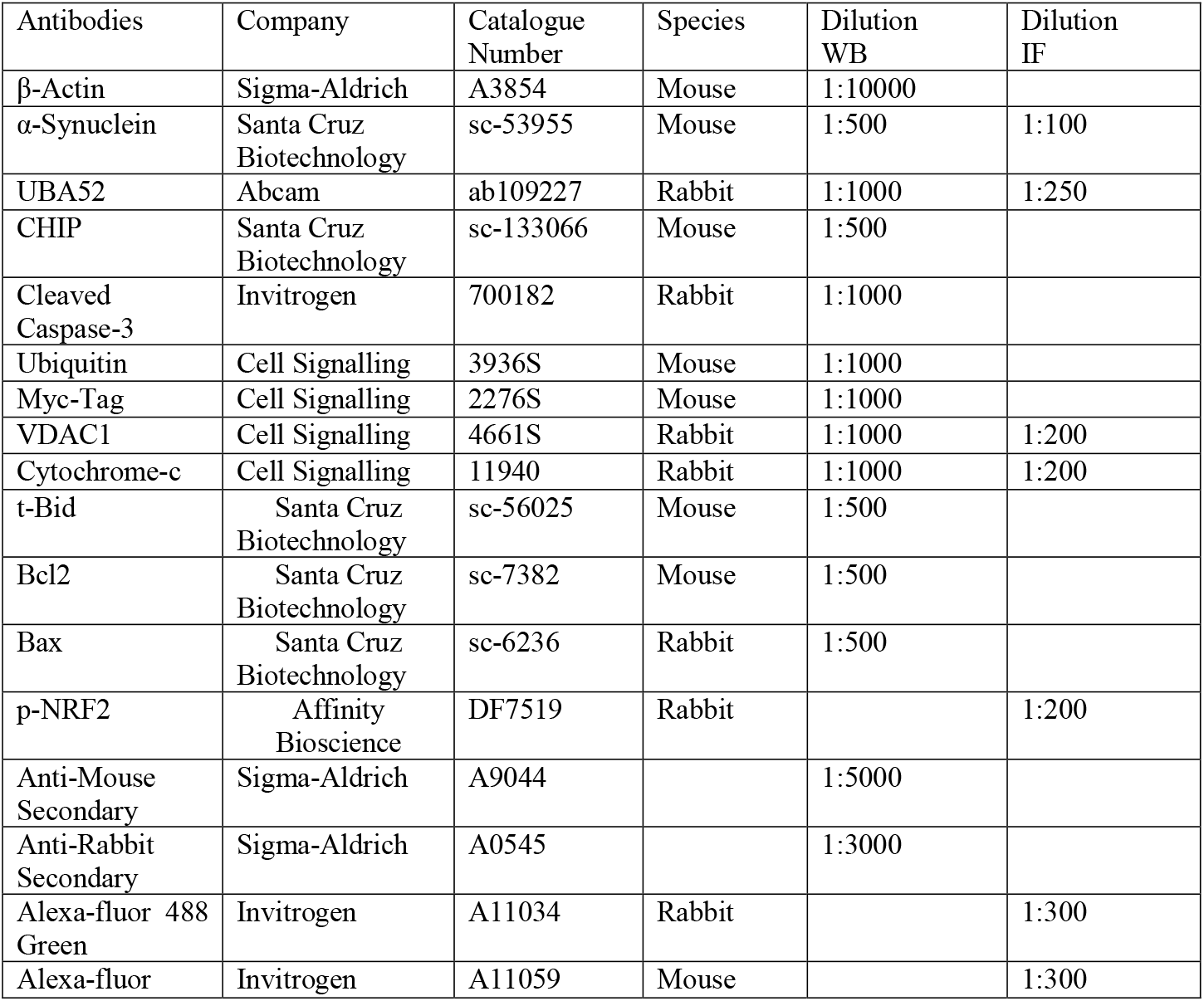
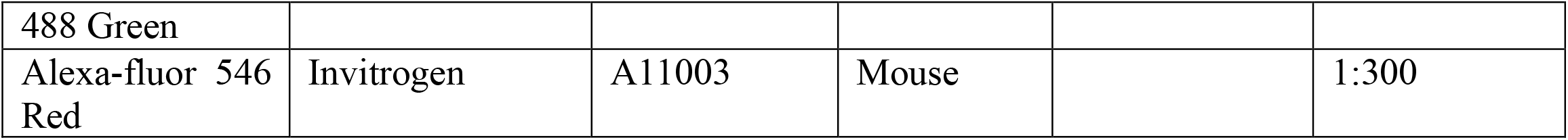
Antibodies used for Immunoblot (WB) and immunofluorescence (IF)

